# Imlifidase and EndoS enables semi-allogeneic bone marrow engraftment in sensitized mice under reduced intensity conditioning

**DOI:** 10.1101/2025.09.10.674224

**Authors:** Marcos I. Petersen, Jiaxin Lin, Kevin Zhan, Perveen Anwar, Robert Bockermann, Colin C. Anderson

## Abstract

Patients in need of a hematopoietic stem cell transplant frequently have pre-existing donor-specific antibodies (DSA) which can impair engraftment. These patients often require intensified conditioning regimens, even in the setting of partially matched (semi-allogeneic) donors. Reducing the toxicity of conditioning and desensitization protocols is therefore a major goal. We previously showed that enzymatic desensitization of donor-specific IgG using imlifidase and EndoS improved murine bone marrow engraftment in donor sensitized, autoimmune-prone recipient mice. Conditioning included a 6 Gy total body irradiation, cyclophosphamide, bortezomib, and T cell depletion. In non-autoimmune prone semi-allogeneic recipients sensitized to the donor, we demonstrate that desensitisation with imlifidase and EndoS permits long-term donor bone marrow engraftment and induces tolerance to subsequent allogeneic skin grafts under a 3 Gy irradiation protocol. Enzymatic inactivation of DSA facilitates donor hematopoietic stem cell engraftment in allo-sensitized semi-allogeneic recipients in a reduced intensity conditioning protocol.

## INTRODUCTION

Pre-existing donor specific antibodies (DSAs) caused by prior sensitization limit engraftment of allogeneic donor bone marrow [1–4]. The presence of DSA is associated with primary graft failure even in the setting of haplo-identical hematopoietic stem cell (HSC) transplantation [5–8]. Current desensitization approaches are varied, complex, and are associated with significant morbidity. There is still a medical need for simpler, more effective, and less toxic approaches [6,9,10].

Imlifidase, an IgG degrading enzyme of *Streptococcus pyogenes* cleaves IgG preventing complement activation and antibody dependent cellular cytotoxicity (ADCC) [11,12]. Imlifidase is conditionally approved in Europe for desensitization of highly sensitized patients before kidney transplantation [13,14]. The successful application of imlifidase in kidney transplantation cannot be directly translated into the field of bone marrow transplantation due to differences between hematopoietic and solid organ transplants. First, except for endothelial cells, solid organs (e.g. kidney) are less sensitive to antibody mediated rejection due to the reduced accessibility of antibodies and complement factors. In contrast, cells of bone marrow transplants circulate in the blood, where complement and donor specific antibodies (DSA) are present, making hematopoietic cells more susceptible to lower levels of DSA [15].

Furthermore, the expression level of donor antigens (MHC/HLA) are expected to be much lower on kidney cells than on hematopoietic cells. In bone marrow transplantation (BMT), the number of cells transplanted is also much lower compared to a kidney. Therefore, when a sensitized patient undergoes BMT, each donor bone marrow cell will be exposed to more DSA than donor kidney cells would be. Thus, the acceptable level of DSA in kidney transplantation is typically higher than in BMT [1,16,17].

To test whether inactivation of IgG with bacterial derived enzymes could facilitate allogeneic bone marrow engraftment in sensitized recipients, we combined imlifidase with Endoglycosidase of *S. pyogenes* (EndoS), an enzyme that de-glycosylates IgG, lowering its binding to FcR and activation of complement [18–22]. We used this combination of enzymes as imlifidase does not cleave IgG1 or IgG2b in mice [21] while it cleaves all four IgG isotypes in humans. We previously demonstrated that Imlifidase together with EndoS prevented bone marrow rejection, allowing long-term fully mismatched allogeneic chimerism in approximately 60 % of recipient mice [23]. In this prior proof of principle study, the donor sensitized recipient mice were completely resistant to chimerism induction when the enzymes were not included, despite an extensive conditioning protocol that included 6 Gy total body irradiation (TBI). The recipients were NOD mice, an autoimmune prone strain resistant to tolerance induction and relatively resistant to irradiation [24,25]. These factors, together with the recognition that hematopoietic stem cell (HSC) transplants in humans are typically done with partially matched donors, led us to test whether imlifidase/EndoS could enable donor bone engraftment in sensitized semi-allogeneic recipients with a reduced intensity conditioning regimen. This project aimed to develop a low-intensity conditioning regimen for DSA-sensitized patients to achieve tolerance via HSC transplantation, thereby permitting subsequent allogeneic organ transplantation.

## METHODS

### Animals

FVB/NJ (H-2^q^; termed FVB), B6 CD45.1, and NOD.B10Sn-H2^b^/J (H-2^b^; termed NOD.H-2^b^) mice were purchased from the Jackson Laboratory (Bar Harbor, ME, USA). BALB/c (H-2^d^) mice were obtained from Charles River Laboratories International (Wilmington, MA, USA). B6.*Rag2p*^*GFP*^ mice [26,27] were kindly provided by Dr. Pamela Fink (University of Washington, Seattle, WA) and backcrossed to B6 CD45.1 mice to generate B6.*Rag2p*^*GFP*^ CD45.1 mice. Recipients were adult FVB x B6. *Rag2p*^*GFP*^ CD45.1 F1 (termed FVB x B6.*Rag2p*^*GFP*^ F1) mice and donors were adult BALB/c x B6.*Rag2p*^*GFP*^ CD45.1 F1 (termed BALB/c x B6.*Rag2p*^*GFP*^ F1) mice. All mice were bred and housed in a specific pathogen-free facility at the University of Alberta. Care and handling of animals followed the guidelines of the Canadian Council on Animal Care. All mice were between 7 to 9 weeks of age at the time of recipient sensitization. The sex of the animals was balanced in both the sensitized and sensitized plus enzyme treatment groups, with an equal number of male and female mice (50% each). The naïve control group included 4 females and 2 males.

### Reagents for desensitization and conditioning

Imlifidase and EndoS were provided by Hansa Biopharma AB (Lund, Sweden) and used with permission. The mAbs specific for CD4 (clone Gk1.5, rat IgG_2b_), CD8α (clone YTS169.4, rat IgG_2b_) and CD90 (clone 30H12, rat IgG_2b_) were purchased from BioXcell. Cyclophosphamide (29875) and bortezomib (A2614) were obtained from Sigma (MO, USA) and ApexBio (TX, USA), respectively.

### Serum DSA detection assay

FVB x B6.*Rag2p*^*GFP*^ F1 mice were sensitized by i.p. administration of 20 × 10^6^ BALB/c x B6.*Rag2p*^*GFP*^ F1 splenocytes. Serum samples were collected prior to sensitization, four weeks post-sensitization, as well as at 4 hours post imlifidase and EndoS treatment. BALB/c x B6.*Rag2p*^*GFP*^ F1 splenocytes (2 × 10^5^) were treated with FcR blockade (anti-mouse CD16/CD32 rat IgG_2b_ antibodies, clone 2.4G2, BE0307, Bio X cell) for 5 minutes, followed by incubation with a titrated quantity of sera in 100μL for 30 minutes. Cells were washed twice and incubated with fluorochrome conjugated secondary antibodies in 100μL for 30 minutes. The following secondary antibodies from Jackson ImmunoResearch were used: FITC conjugated F(ab’)_2_ fragment from rabbit anti-mouse IgG Fc antibody (315-096-046), and AF647 conjugated goat anti-mouse IgG Fc antibody (115-606-071). Cells were washed twice and analyzed by flow cytometry. HBSS with 2% FBS was used for cell washes and reconstitution.

### BMT protocol and monitoring of chimerism

Naïve or donor sensitized FVB x B6.*Rag2p*^*GFP*^ F1 (CD45.1) mice received a donor BALB/c x B6.*Rag2p*^*GFP*^ F1 BMT; sensitization was by i.p. injection of 20 × 10^6^ donor (F1) splenocytes between 29-42 days prior to BMT; a booster immunization was given at 21 days prior to BMT to three mice with low DSA (2 in the sensitized group and 1 in the sensitized plus enzymes group). All recipients were given the following conditioning regimen with or without 100 μg imlifidase and 30 μg EndoS i.v. 4 hours before BMT. Conditioning: Cyclophosphamide (150 mg/kg, i.p.) and bortezomib (1 mg/kg, i.p.), referred to as CyBor, were given on day −4. T cell-depleting antibodies were administered i.p. on day −6, −2, 2, 6, 11, and 16. Four hours prior to BMT all recipients received 3 Gy of TBI (Gammacell 3000). Donor bone marrow cells (80 × 10^6^) were given intravenously (i.v.) via the lateral tail vein on day 0. Flow cytometry analysis on peripheral blood at the time points indicated was done to monitor chimerism. Donor mice were CD45.1/2 heterozygous, allowing discrimination from recipient cells by differential CD45 allelic expression using anti-CD45.2 antibody.

### Antibodies and flow cytometry

Fluorochrome-labeled antibodies against mouse CD45.2 (104), B220 (RA3-6B2), TCRβ (H57-597), CD4 (RM4-4), CD8β (H35-17.2), CD11b (M1/70), CD11c (N418), NK1.1 (PK136), Ly6G (1A8) were purchased from BD Pharmingen (CA, USA) or Tonbo Biosciences (CA, USA). An LSR II (Becton Dickson, CA, USA) flow cytometer was used for data acquisition, and data analysis was performed using FlowJo (Treestar software, OR, USA).

### Skin grafting

Four months post BMT, some of the recipients were given skin grafts. Approximately 1 cm^2^ full thickness trunk skin from sex-matched donor BALB/c x B6.*Rag2p*^*GFP*^ and 3^rd^ party NOD.H-2^b^ were transplanted onto the graft beds created on the dorsum of recipients, with approximately 1 cm distance in between grafts. Skin grafts were secured with sutures and bandages were maintained for seven days. The grafts were inspected weekly after removal of the bandage and scored as rejected when approximately 90% or more of the surface area had been eliminated or had become necrotic.

### Statistical analysis

All statistical analyses were conducted using R version 4.4.1 (R Foundation for Statistical Computing, Vienna, Austria). Chimerism frequencies at day 112 were compared across three experimental groups: sensitized, sensitized plus enzyme-treated, and naïve controls. Pairwise comparisons were performed by constructing 2 × 2 contingency tables representing the presence or absence of chimerism in each group. Differences in chimerism rates were evaluated using two-sided Fisher’s exact tests. A P value less than 0.05 was considered statistically significant. Donor skin graft survival was analyzed using the Kaplan– Meier method, and differences between groups were assessed with the log-rank (Mantel-Cox) test using the survival package in R [28].

## RESULTS AND DISCUSSION

### Imlifidase and EndoS facilitate semi-allogeneic BMT and skin graft acceptance in recipients with DSA

To examine whether the enzymes could facilitate chimerism in sensitized semi-allogeneic recipients given a reduced intensity conditioning protocol (Fig. 1), we used F1 donor and recipient mice, both having a *Rag2p*^*GFP*^ transgene. This GFP reporter model allows the tracking of newly generated B cells and T cells that were recent thymic emigrants (RTE) [26,27]. The donor recipient combination included a 1 haplotype mismatch and multiple minor antigen mismatches. To generate donor sensitized recipients, we immunized FVB x B6.*Rag2p*^*GFP*^ F1 mice with donor cells (BALB/c x B6.*Rag2p*^*GFP*^ F1) and measured the level of DSA by flow cytometry [23]. Sensitized recipients with DSA ≥ 5,000 MFI and naïve (not sensitized) recipients were subsequently used for BMT studies. The DSA 5000 MFI cutoff was set based on a pilot study where some recipients with levels below 4000 MFI became chimeric, indicating insufficient sensitization. All naïve recipients given the conditioning protocol without enzyme treatment achieved a high level of donor cell chimerism that was maintained throughout the observation period (more than 100 days; Fig. 2). None of the sensitized recipients achieved stable long-term chimerism without enzyme treatment. However, one out of six of these recipients, that had a moderate level of DSA, did have transient chimerism. In contrast, six out of eight sensitized and enzyme treated recipients maintained a high level of chimerism for more than 100 days (chimerism at day 112 was 0/6 for sensitized and 6/8 for sensitized enzyme treated groups, *P* = 0.01 Fisher’s exact test). Chimerism was unstable in 2 sensitized enzyme treated recipients (SE1 and SE3, Fig. 2D), eventually leading to a complete loss of chimerism; not surprisingly, unstable chimerism was associated with recovery of high levels of recipient T cells in both mice and one of the unstable chimeras had a very high level of DSA (Fig. 2 E, F). Long-term stable donor cell chimerism in sensitized enzyme treated recipients included multiple lineages of donor cells, including T cells, B cells, NK cells, neutrophils, dendritic cells and macrophages (Fig. 2). The development of high levels of donor T cell chimerism occurred earlier, by day 84, for naïve recipients compared to the sensitized enzyme treated group; even by day 112 a substantial portion of T cells in most mice of the sensitized enzyme treated group were of recipient origin (Fig. 3). Optimal immune reconstitution with donor hematopoietic cells would be expected to include newly generated donor T and B cells and not just homeostatic expansion of mature cells present in the donor BMT. Thus, we expected to find that a portion of donor T and B cells in chimeras would transiently express GFP, indicating they had recently expressed the *Rag2* gene (i.e. were newly generated cells); other lineages would not be expected to express GFP. In the recipients with high levels of stable multilineage chimerism, for both the naïve and sensitized enzyme treated groups, the donor chimerism included a substantial portion of recent thymic emigrants and newly generated B cells; the absence of GFP in NK cells is shown as a negative control (Fig. 3). Thus, the conditioning regimen and semi-allogeneic BMT did not compromise thymus function, and donor hematopoiesis, including T and B cell development, was maintained long-term. The semi-allogeneic BMT, using this conditioning protocol, did not lead to any overt graft versus host disease, as there was no loss of weight in BMT recipients (all gained weight compared to pre-transplant) and they appeared healthy throughout the experiment.

**Figure 1.**
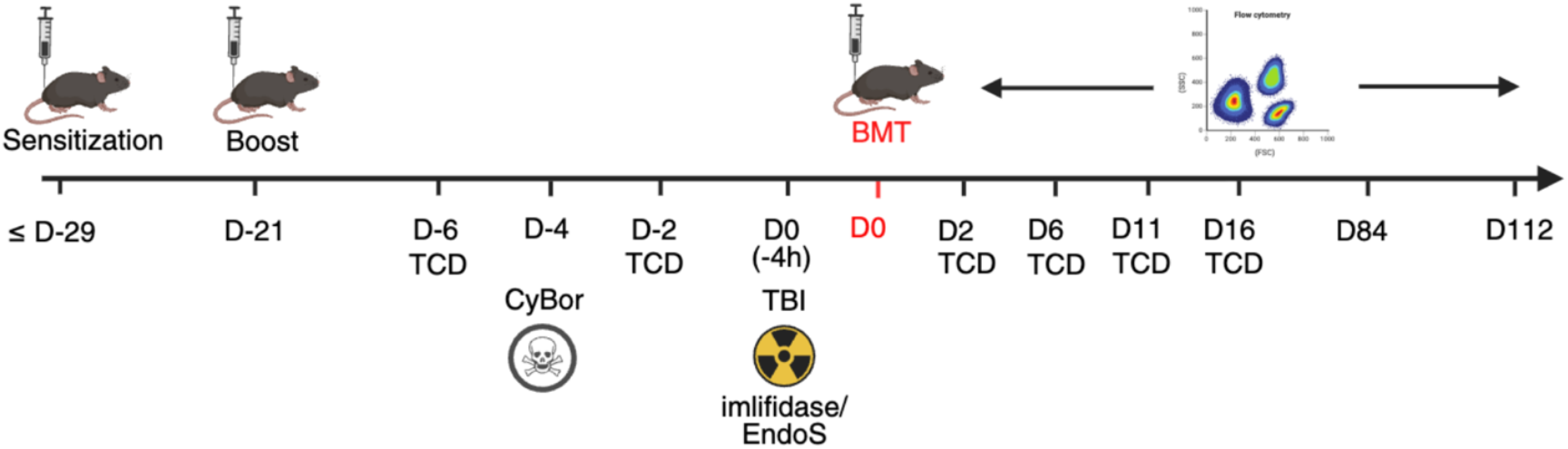
Reduced intensity conditioning protocol for semi-allogeneic bone marrow transplantation in sensitized recipients. Timing, in days (D) relative to bone marrow transplantation (BMT). Mice were sensitized between D-42 and D-29. Individuals with low DSA levels received a booster at D-21. Treatment with antibodies to achieve T cell depletion (TCD), total body irradiation (TBI), enzyme (imlifidase/EndoS) administration, and flow cytometry on peripheral blood cells, is depicted. Created in BioRender. Bockermann, R. (2025) https://BioRender.com/ydnm7zb

**Figure 2.**
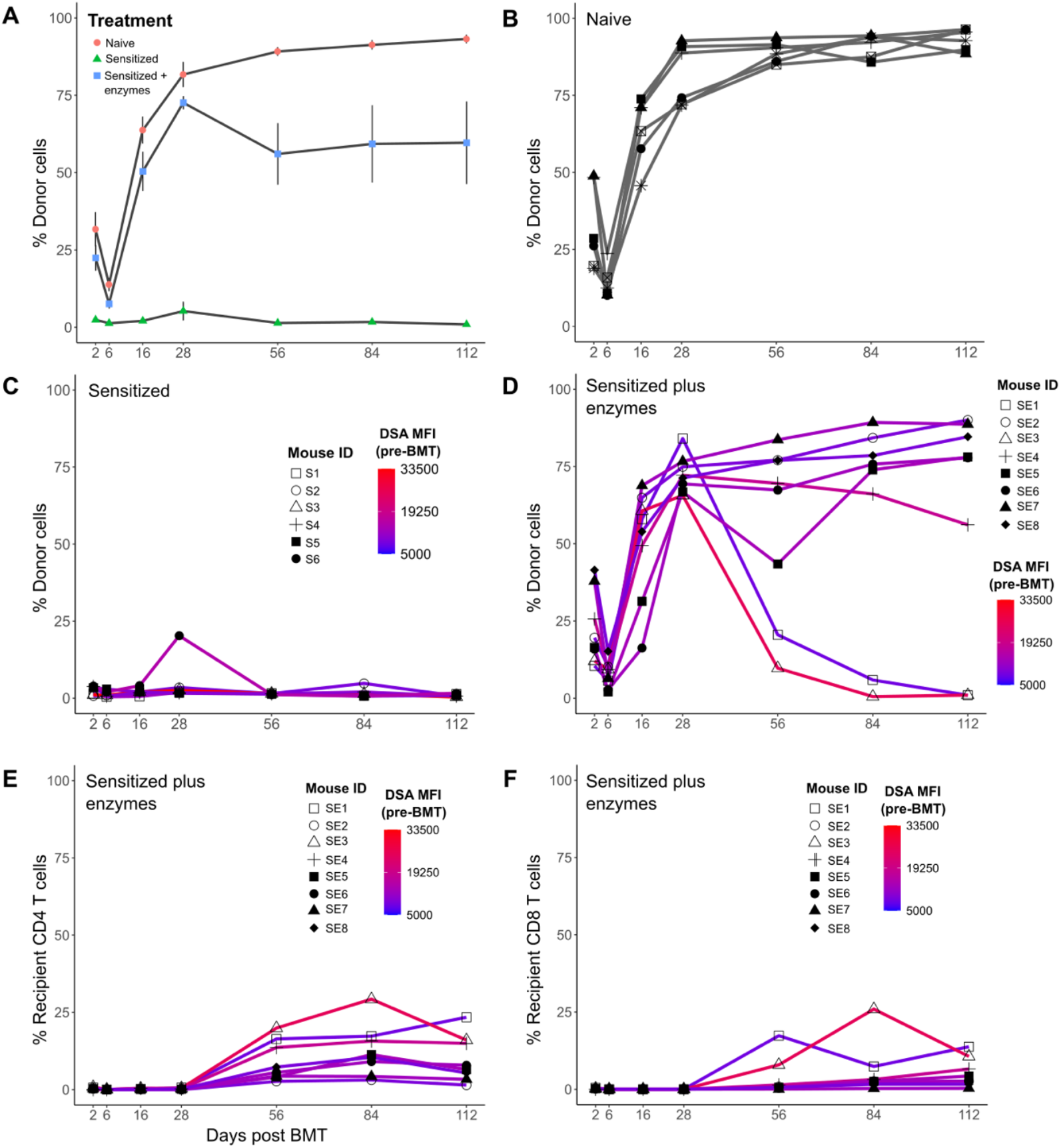
Imlifidase plus EndoS prevent rejection of semi-allogeneic bone marrow cells in sensitized recipients under a reduced intensity conditioning regimen. For the indicated groups, donor and recipient cells in peripheral blood were assessed by flow cytometry and the proportion of donor cells of all cells are shown as the mean ± S.E.M. (A) and for individual mice (B-D). The proportion of all cells that are recipient CD4 T cells (E) and CD8 T cells (F) are shown for individual mice of the sensitized plus enzymes group. The level (MFI) of DSA for each recipient is depicted based on the colour scale shown (C-F). DSA was determined 6 days prior to BMT.

**Figure 3.**
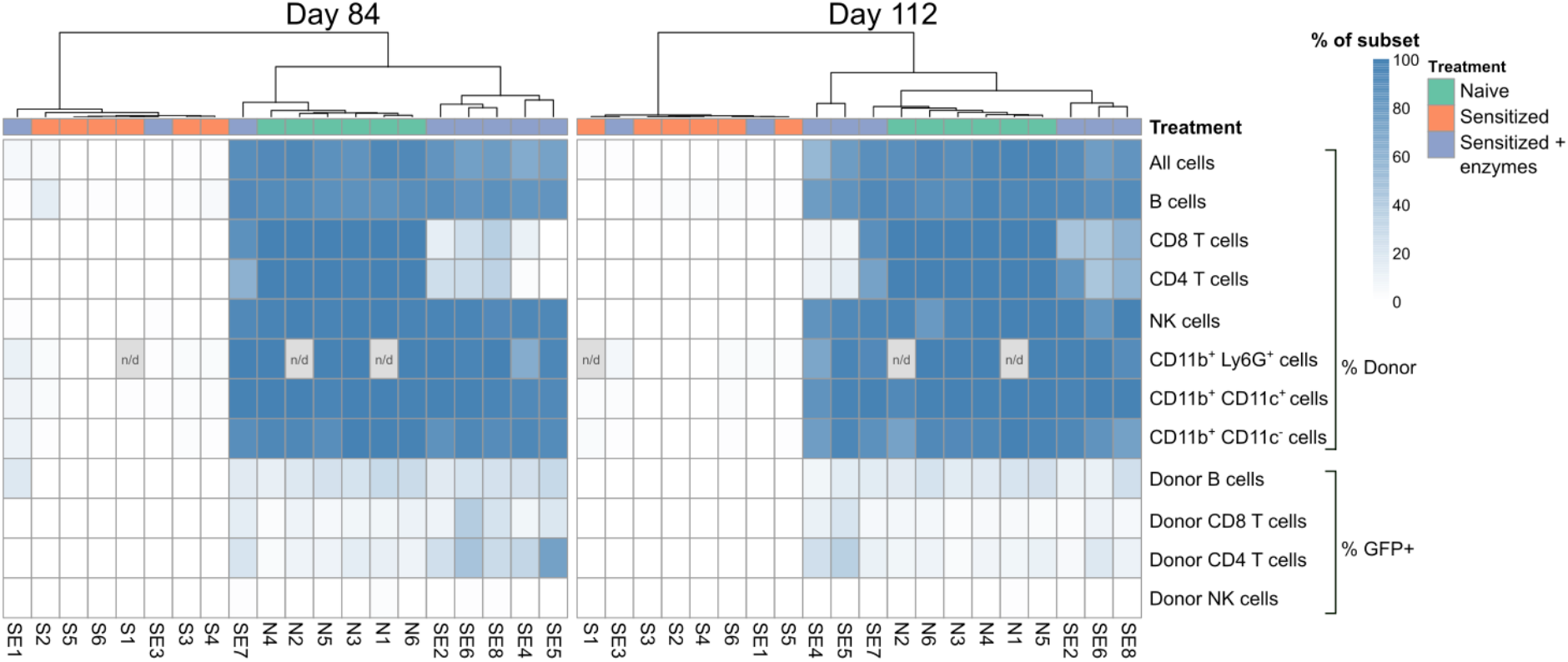
Chimerism in sensitized recipients treated with imlifidase and EndoS is multilineage and includes newly generated donor T and B cells. Hierarchical clustering of individual recipient peripheral blood samples, on day 84 (left) and day 112 (right), based on the proportion of donor cells of the cell lineages shown and the proportion of donor T, B and NK cells expressing GFP (GFP is expressed in newly generated T or B cells), depicted as a heatmap. Individual mouse ID’s are depicted along the bottom and correspond to mouse ID’s shown in Fig. 1. (S = sensitized; SE = sensitized plus enzymes; N = non-sensitized), n/d = not determined for the indicated sample.

In some cases, bone marrow transplantation can lead to a state known as split tolerance, where the recipient is tolerant of some donor cells or tissues but not others. For example, a state of chimerism with donor T cells without other donor hematopoietic lineages does not typically allow acceptance of a donor skin graft [29]. The presence of multilineage chimerism in sensitized enzyme treated recipients suggested they may be tolerant of a wide array of donor antigens. To test this possibility and to test whether these mice maintained immunocompetence, we gave a cohort of each recipient group a donor and 3^rd^ party skin graft. The 3^rd^ party graft (NOD.H-2^b^ donor) was mismatched only for minor histocompatibility antigens to provide a more stringent test of immunocompetence. As shown in Table 1, all sensitized recipients without enzyme treatment rejected both donor and 3^rd^ party skin grafts. In contrast, of five sensitized enzyme treated recipients, four rejected the 3^rd^ party graft and only one rejected the donor skin graft. Not unexpectedly, the one recipient that rejected its donor skin graft was a recipient with very high DSA and transient chimerism (mouse SE3, chimerism data shown in Fig. 2). Thus, most sensitized enzyme treated recipients achieved stable chimerism and a tolerance that extended to donor skin grafts. In addition, the majority maintained and/or recovered sufficient immunocompetence to reject a multiple minor histocompatibility mismatched skin graft. While we did not examine DSA levels post BMT, in most stable long-term chimeras the vast majority (> 80 %) of B cells in circulation were of donor origin and therefore a rebound in DSA would not be expected. Altogether, these data provide pre-clinical data indicating that IgG degrading enzymes provide a successful desensitization approach to prevent rejection of a semi-allogeneic bone marrow transplant in sensitized recipients given a reduced intensity conditioning regimen. It is noteworthy that a single dose of enzymes was sufficient to prevent bone marrow rejection in most recipients; multiple doses or a single higher dose might be required for those with the highest levels of DSA. Overall, these pre-clinical data support the use of a simplified desensitization protocol for allogeneic bone marrow transplant recipients using enzymes that temporarily degrade the function of recipient IgG.

**Table 1.** Donor skin graft acceptance by sensitized recipients depends on enzyme mediated desensitization (skin graft survival, days)

| Recipient | Donor skin graft | 3<sup>rd</sup> Party skin graft | significance |
| --- | --- | --- | --- |
| Naive | > 100 x 3 | 14, 30, n/d |  |
| Sensitized | 14, 21 x 2, 28 | 14, 21 x 3 |  |
| Sensitized plus enzymes | <sup>#</sup>14, > 100 x 4 | 20, 28 x 3, > 100 | <sup>*</sup>*P* = 0.04<br><sup>‡</sup>*P* = 0.04 |

Survival time of skin grafts in days. Two skin grafts, donor (BALB/c x B6.*Rag2p<sup>GFP</sup>* F1) and a 3<sup>rd</sup> party (NOD.H-2<sup>b</sup>), were given to FVB x B6.*Rag2p<sup>GFP</sup>* F1 recipient mice 5-6 months post BMT. n/d = not determined (graft technical failure).
<sup>#</sup>The mouse that rejected the donor skin graft at day 14 had lost chimerism prior to skin grafting (see mouse SE3 in Fig. 1).
<sup>\*</sup>Log rank test *P* value comparing donor skin graft survival between the sensitized and the sensitized plus enzymes groups.
<sup>‡</sup>Log rank test *P* value comparing donor vs. 3<sup>rd</sup> party skin graft survival in the sensitized plus enzymes group.

## Abbreviations

BMC: bone marrow cells
BMT: bone marrow transplantation
CyBor: bortezomib and cyclophosphamide
DSA: donor specific antibodies
EndoS: endoglycosidase of *Streptococcus pyogenes*
GVHD: graft-versus-host disease
IdeS: IgG degrading enzyme of *Streptococcus pyogenes*
IVIG: intravenous immune globulins
TBI: total body irradiation
HSC: hematopoietic stem cell

## DECLARATION OF GENERATIVE AI AND AI-ASSISTED TECHNOLOGIES IN THE WRITING PROCESS

AI related technologies were not used in the writing of this manuscript.

## ACKNOWLEDGMENTS

We thank HSLAS staff for assistance with animal care. This work was supported by Hansa Biopharma. *Authorship Contributions:* J.L. and M.I.P. designed and performed experiments, analyzed data, and wrote the manuscript; C.C.A. designed experiments, analyzed data, and wrote the manuscript; P.A. performed experiments and revised the manuscript; R.B. designed experiments, interpreted data and revised the manuscript.

